# SPACA9 Acts as a Molecular Staple Modulating Microtubule Dynamic Instability

**DOI:** 10.64898/2026.06.14.732105

**Authors:** Mohammed Aboraya, Shulamit Fluss Ben-Uliel, Ron Orbach

## Abstract

Motile cilia rely on highly stable axonemal microtubules reinforced by microtubule inner proteins (MIPs) that form a network within their lumen, yet the functions of individual MIPs remain poorly understood. Here, we characterize the conserved MIP sperm acrosome-associated protein 9 (SPACA9), which localizes to respiratory cilia and sperm flagella. Using *in vitro* reconstitution assays, we show that human SPACA9 (hSPACA9) acts as a molecular staple: it stabilizes protofilaments at growing microtubule ends, and inhibits dynamic instability. Surprisingly, these effects do not confer resistance to motor-induced lattice damage, indicating that regulation of microtubule dynamics can be uncoupled from mechanical resilience. Mechanistically, we identify an unstructured C-terminal region that is sufficient for microtubule binding and recapitulates the effects on dynamics. Together, our findings reveal functional specialization among MIPs and provide a mechanistic framework for how lumenal proteins tune distinct properties of axonemal microtubules.

## Introduction

Eukaryotic motile cilia and flagella are present in nearly all eukaryotes and are used for locomotion as well as generating fluid flow (1). The structural backbone of cilia is the axoneme, a highly stable microtubule-based structure (2). In most motile cilia, the axoneme consists of nine outer doublet microtubules (DMTs) surrounding a central pair of singlet microtubules, forming the stereotypical 9+2 architecture (3–5). Each DMT is composed of a complete 13-protofilaments A-tubule and an incomplete 10-protofilaments B-tubule (6). The axoneme serves as a scaffold for numerous proteins and protein complexes that associate with the external surface of the ciliary microtubules, such as the radial spokes and dynein arms (7, 8). Additionally, dozens of proteins, collectively known as microtubule inner proteins (MIPs), form an interwoven network within the ciliary microtubule lumen (4, 9, 10).

The axoneme provides structural support for cilia while also withstanding mechanical demands during ciliary beating and protein trafficking. During beating, cyclic bending generates strain energy in the microtubule lattice, potentially promoting structural destabilization. This mechanical stress is further amplified by motor protein activity, such as that of dynein arms, and the forces imposed by intraflagellar transport (IFT) (11). In addition to these externally applied forces, microtubules experience intrinsic mechanical stress arising from the tubulin GTPase cycle. The curved structure of the tubulin dimer introduces strain into the polymerized microtubule lattice, which increases following GTP hydrolysis. Weakened lateral contacts may reduce the resistance of the lattice to this strain, destabilizing the lattice, and favoring shrinkage onset (12–14). As a result, microtubules undergo dynamic instability, a process in which they alternate between phases of growth and shrinkage through transitions known as catastrophe (growth to shrinkage) and rescue (shrinkage to growth) (15, 16).

In the cilia, where mechanical stress and hydrolysis-induced strain energy are ubiquitous, stabilization mechanisms are essential to suppress microtubule dynamic instability and maintain structural integrity. Thus, the complex network of MIPs has been suggested to provide a mechanical framework. This structural reinforcement is thought to increase microtubule rigidity and resilience, enabling the axoneme to withstand mechanical stress during ciliary beating (4, 17–19). However, the interwoven network of MIPs within the ciliary microtubule lumen, combined with functional redundancy, complicates the determination of their specific contributions, as the knockout of a specific MIP often leads to the loss of others (18, 20–24). Thus, the necessity for such a large number of MIPs, as well as the underlying mechanisms of function and specific roles of each protein, remain poorly understood.

The sperm acrosome-associated 9 (SPACA9) protein is an evolutionarily conserved MIP localized within the cilia of the respiratory tract and the flagella of sperm cells across metazoans (25–29). In the B-tubule of DMTs, SPACA9 forms periodic intralumenal striations with an 8-nm repeat, aligning along protofilaments. In singlet microtubules extending from the doublet at the flagellar endpiece (30, 31), SPACA9 organizes into a discontinuous spiral structure known as TAILS (Tail Axoneme Intralumenal Spiral), which also exhibits an 8-nm periodicity. It was further shown that SPACA9 engages the microtubule lumen through a structured α-helical core; however, the C-terminal tail is not resolved in the structure, leaving its contribution to microtubule interaction unknown. Thus, despite its conservation and defined localization, the functional contribution of SPACA9 to ciliary microtubules has yet to be elucidated.

To begin dissecting how individual MIPs contribute to axonemal stability, we focused on the conserved protein human SPACA9 (hSPACA9) as a model. Using a reductionist *in vitro* approach, we reconstituted hSPACA9 with microtubules to examine its binding behavior and its effects on microtubule dynamic instability. Through truncation and mutational analysis, we further probed the contribution of distinct regions of the protein, including the previously uncharacterized C-terminal tail. Finally, we tested whether hSPACA9 can protect microtubules from motor-induced mechanical stress using motility-based assays.

## Results

### hSPACA9 binds microtubules and GFP tagging impairs its functional interaction

To investigate the effect of hSPACA9 on microtubules, we expressed and purified recombinant hSPACA9^GFP^ by two chromatography steps (Fig. S1A). While the GFP tag enables direct visualization of protein-microtubule interactions, we hypothesized that its size, comparable to that of hSPACA9, could interfere with lattice association due to steric constraints in the lumen. To address this possibility, we also expressed and purified unlabeled recombinant hSPACA9 using the same procedure (Fig. S1B). Circular dichroism (CD) spectroscopy was used to assess the secondary structure of hSPACA9. The far-UV CD spectrum displayed pronounced minima at approximately 208 and 222 nm, characteristic of a predominantly α-helical fold (Fig. S1C). Thus, the CD data are consistent with the previously resolved structure of hSPACA9, which reveals a predominantly α-helical architecture.

To assess whether the two hSPACA9 constructs interact with microtubules *in vitro*, we polymerized microtubules in the presence of increasing concentrations of either hSPACA9 or hSPACA9^GFP^ and conducted co-sedimentation assays. In both cases, hSPACA9 and hSPACA9^GFP^ were detected in the pellet fraction, confirming that both constructs of hSPACA9 interact with microtubules (Fig. 1A). In contrast, no binding of hSPACA9 to free tubulin was observed in a pull-down assay (Fig. S2). Since many microtubule-associated proteins bind via the negatively charged C-terminal E-hooks, which mediate interactions on the microtubule surface, we examined whether hSPACA9 interacts nonspecifically with the external lattice. Microtubules were treated with subtilisin to remove the E-hooks, however, hSPACA9-GFP fluorescence was still observed on the microtubules after treatment (Fig. S3).

**Figure 1.**
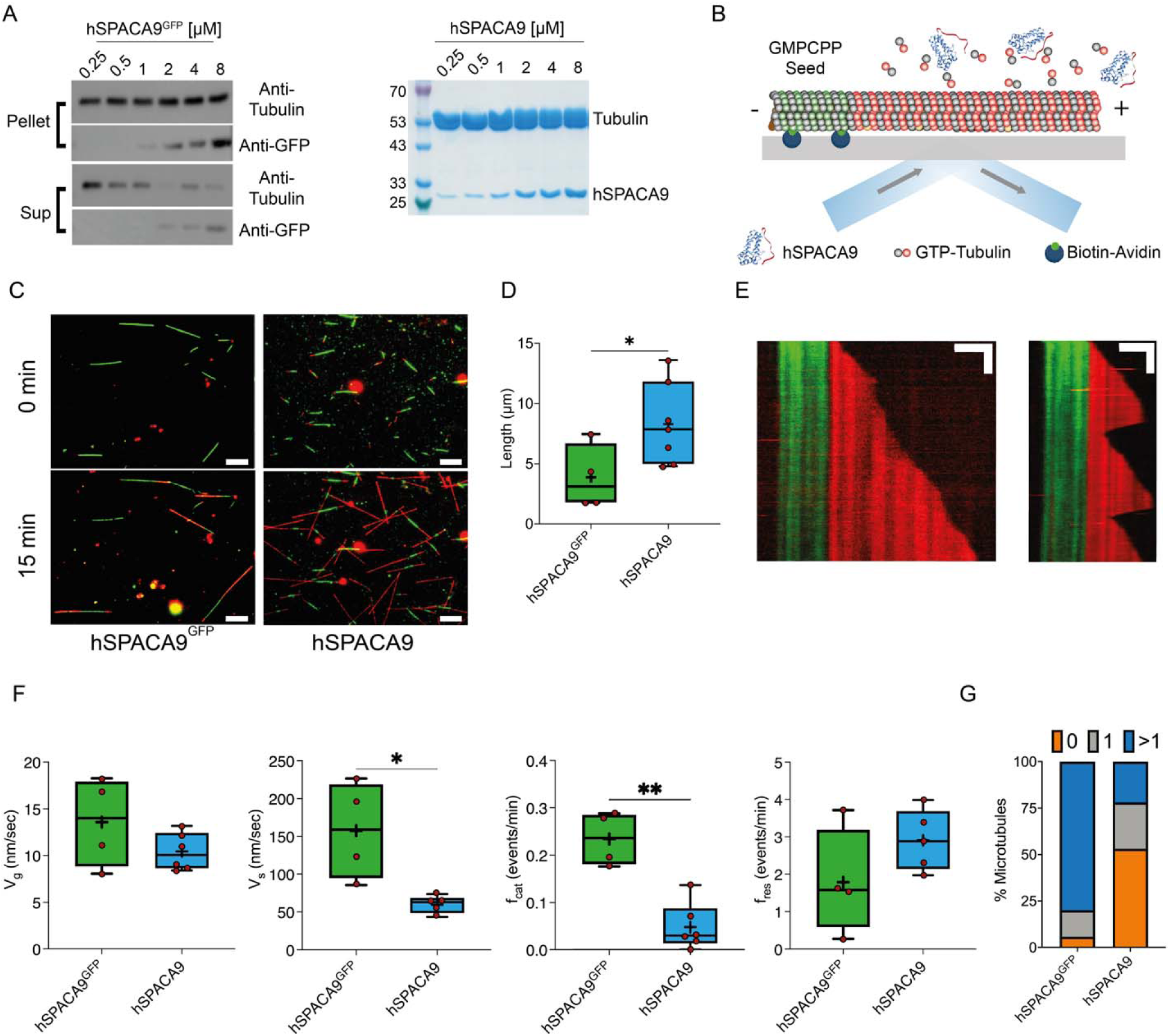
hSPACA9 binds to microtubules *in vitro*. (**A**) Western blot analysis of co-sedimentation assay in the presence of hSPACA9^GFP^ (left), and Coomassie-stained SDS-PAGE of microtubule pellet fractions following co-sedimentation assay with hSPACA9 (right). (**B**) Schematic representation of TIRFM *in vitro* assay. (**C**) Representative TIRFM images of dynamic microtubules in the presence of 300nM hSPACA9^GFP^ and hSPACA9 at time 0 and after 15 min. Orange extensions indicate regions where hSPACA9^GFP^ binds to the red dynamic microtubules. Scale bar: 5 µm. (**D**) Tukey plot showing the mean extension length. Horizontal line within each box represents the median, and the plus sign indicates the mean. Each red circle marks the mean growth rate of a single biological replicate, n =188, 226; data pooled across at least 4 independent experiments. (**E**) Representative kymographs of dynamic microtubules growing from GMPCPP-stabilized seeds (green) in the presence of 10 µM TAMRA-labeled tubulin (red) and 300 nM unlabeled hSPACA9. Scale bars: 2 μm (horizontal), 2 min (vertical). (**F**) Tukey plots illustrating of microtubule plus-end dynamic properties in the presence of hSPACA9^GFP^ or hSPACA9: growth rate (V_g_), shrinkage rate (V_S_), catastrophe frequency (f_cat_) and rescue frequency (f_res_). Each red circle marks the mean growth rate of a single biological replicate. Data pooled across at least 4 independent experiments. All experiments were performed in the presence of 10 μM tubulin, with 300 nM hSPACA9^GFP^ or hSPACA9. *P< 0.05; **P< 0.01 as determined by Mann-Whitney test. In all Tukey plots, the horizontal line within each box represents the median, and the plus sign indicates the mean. (**G**) Bar plot showing the fraction of the microtubule population exhibiting no catastrophe event (orange), a single catastrophe (gray), or multiple catastrophe events (blue), n =34, 95.

### hSPACA9 suppresses microtubule dynamic instability

To further examine whether the two constructs influence microtubule dynamic properties, we performed *in vitro* dynamic assays using total internal reflection fluorescence microscopy (TIRFM) (Fig. 1B) (32, 33). Microtubules were polymerized from guanosine-5’-[(α,β)-methyleno]triphosphate (GMPCPP)-stabilized seeds in the presence of either hSPACA9 or hSPACA9^GFP^. Interestingly, after 15 minutes, microtubule extension lengths in the presence of hSPACA9 were significantly longer compared to those in the presence of hSPACA9^GFP^ (8.3 ± 1.27 μm vs 3.87 ± 1.35 μm, respectively, mean ± SE) (Fig. 1C and D, movies S1–S3). We performed kymograph analysis to examine the microtubule plus-end dynamics (Fig. 1E). The analysis revealed that this increase in extension length for the untagged construct did not result from a significant difference in microtubule growth rate (10.42 ± 0.81 nm/sec vs. 13.57 ± 2.4 nm/sec). Instead, this effect is attributed to significant decrease in shrinkage rate (59.28 ± 5.1 nm/sec vs.157.4 ± 32.53 nm/sec) and catastrophe frequency (0.05 ± 0.02 events/min vs. 0.23 ± 0.03 events/min), along with a moderate increase in rescue frequency among microtubules that did undergo catastrophe (2.9 ± 0.36 events/min vs. 1.78 ± 0.71 events/min) (Fig. 1F). Additionally, the percentage of microtubules that showed no catastrophe events was significantly higher in the presence of hSPACA9 compared to hSPACA9^GFP^ (Fig. 1G). Thus, while both constructs bind to microtubules, GFP labeling appears to interfere with the binding of hSPACA9 to the microtubule lattice, possibly due to spatial constraints within the microtubule lumen. We therefore proceeded with unlabeled hSPACA9 constructs for our analyses, using GFP-tagged constructs solely to examine physical interactions.

To quantify the effect of hSPACA9 on microtubule dynamics, we systematically examined its effects across a range of concentrations. Microtubule growth rates in the presence of increasing concentrations of hSPACA9 did not change significantly compared to the control lacking hSPACA9 (Fig. 2A). In contrast, hSPACA9 markedly decreased microtubule shrinkage rate by up to 2.2-fold (Fig. 2B). Additionally, we observed a significant decrease in catastrophe frequency (Fig. 2C) and increase in the rescue frequency in the presence of hSPACA9 (Fig. 2D). This modest increase in rescue events, however, is due to the significant decrease in the number of microtubules undergoing catastrophe events (Fig. 2E). Evidently, after normalization of the number of rescue events to the total shrinkage length, a clear increase in the frequency of rescues per unit of shrinkage was detected (Fig. 2F). Thus, hSPACA9 reduces microtubule dynamic instability by shifting the balance toward more persistent growth and reduced transition into depolymerization.

**Figure 2.**
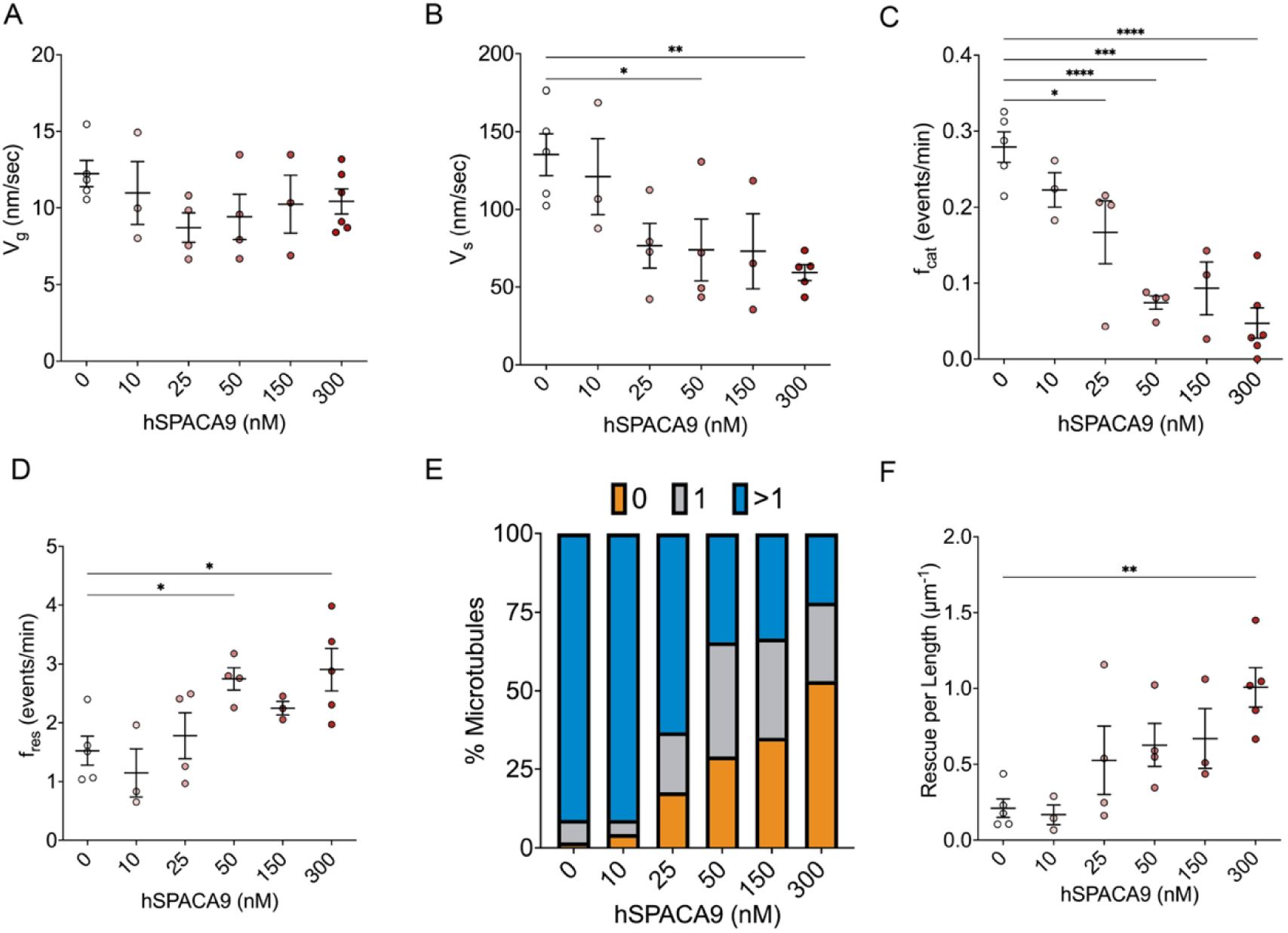
Quantification of microtubule dynamic properties. Dot plots showing the effect of hSPACA9 concentration on (**A**) Microtubule growth rate (V_g_), (**B**) Microtubule shrinkage rate (V_s_), (**C**) Catastrophe frequency (f_cat_), (**D**) Rescue frequency (f_res_). (**E**) Frequency plot showing the percentage of microtubules undergoing varying numbers of catastrophe events at different hSPACA9 concentrations. (**F**) Rescue frequency normalized to microtubule length as a function of hSPACA9 concentration. Each dot represents the mean of a biological replicate, with the red color gradient indicating the corresponding hSPACA9 concentration. Black lines show the overall mean and SE across replicates. All experiments were performed in the presence of 10 µM free tubulin; *P< 0.05; **P< 0.01; ***P< 0.001 ;****P< 0.0001 as determined by one-way ANOVA followed by Dunnett’s multiple comparisons test. Statistical analysis was performed using the average value from each independent experiment; n=45–96.

### hSPACA9 stabilizes protofilaments and promotes curved microtubule tip structures

Previous studies have reported that the microtubule tip region can extend up to ∼200 nm (34, 35). We found that in the presence of hSPACA9, a subset of growing microtubules exhibited curved tip structures characterized by reduced fluorescence intensity, features that were not observed in its absence. These structures gradually straightened as additional protofilaments polymerized along the microtubule shaft (Fig. 3A, movie S4). A subset of microtubules displayed a distinct pause in growth, which was subsequently followed by accelerated elongation (Fig. 3B, movie S5). At high hSPACA9 concentrations (2 μM), spontaneous nucleation was observed, generating microtubules with similar curved tip structures at their growing ends. As in the previously observed cases, these structures gradually straightened following protofilament elongation (Fig. 3C, movie S6). Collectively, the observation of curved tip structures (Fig. S4A), diminished fluorescence intensity (Fig. 3D), and the subsequent straightening of these tips upon continued protofilament elongation suggest that hSPACA9 stabilizes protofilaments. Indeed, transmission electron microscopy (TEM) further supported the TIRFM observations by revealing open microtubule sheets at the tip in the presence of hSPACA9 (Fig. 3E).

**Figure 3.**
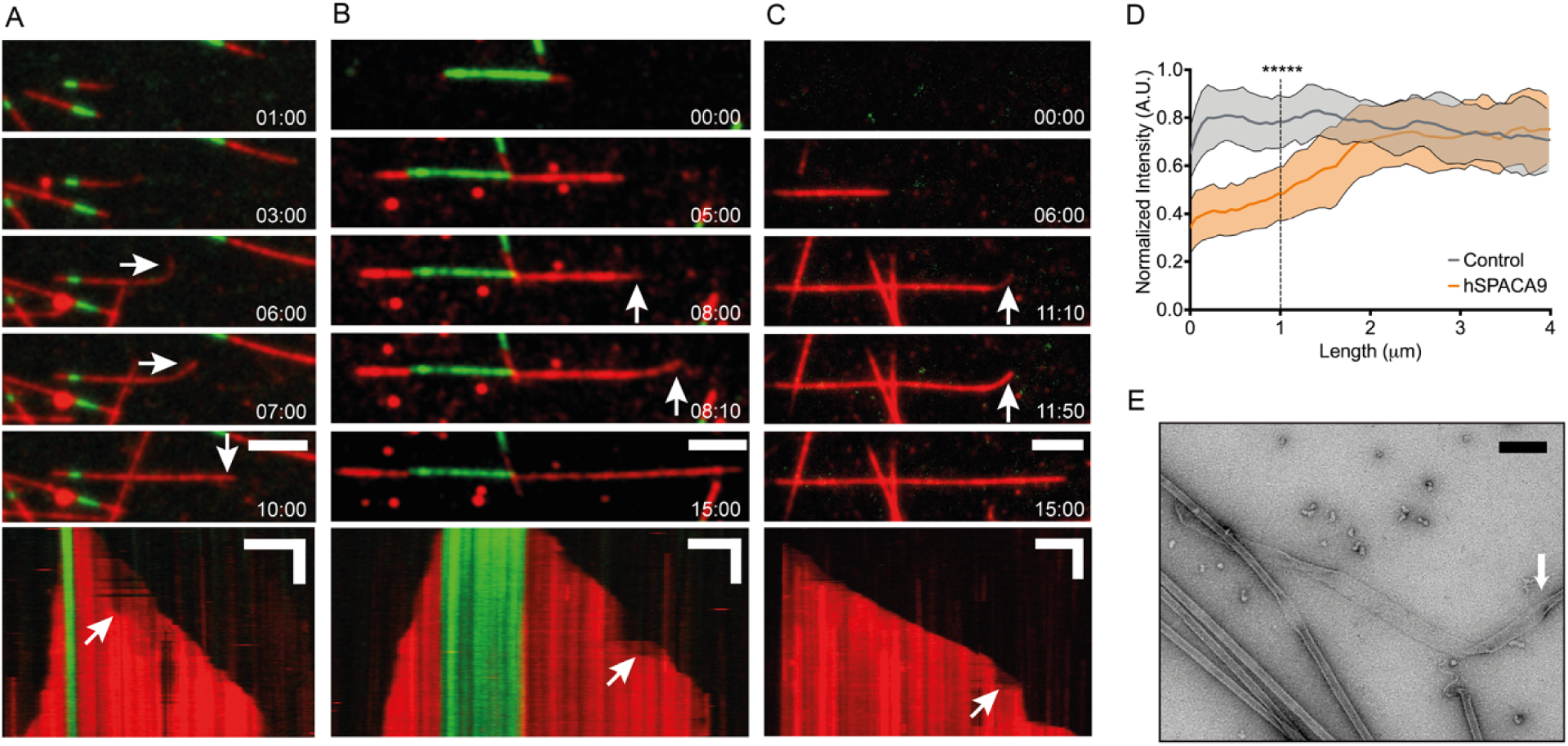
hSPACA9 stabilizes microtubule protofilaments. (**A**) Time-lapse images capture the formation of a curved tip structure in the presence of 300 nM hSPACA9. The kymograph reveals additional growth along the microtubule shaft. (**B**) Time-lapse images showing fast growth of protofilaments at the tip in the presence of 300 nM hSPACA9. (**C**) At high hSPACA9 concentrations (2 µM), spontaneous microtubule nucleation is observed, followed by the formation of a curved tip structure that straightens as additional protofilaments polymerize. Scale bars: 3 μm, 3 min. (**D**) Average normalized intensity profiles of microtubule tip structures in the presence (orange) and absence (gray) of 300nM hSPACA9 reveal reduced fluorescence intensity at the tip region. Error bars represent SD. Statistical significance was assessed at 1LJµm from the tip end, corresponding to the region where all structures remained curved, using an unpaired t-test, n = 24, 32. (**E**) TEM image showing a sheet-like structure at the growing tip of a microtubule (white arrow). Scale bar: 200 nm.

To evaluate the effect on microtubule assembly across a range of tubulin concentrations, we complemented our microscopy assays with a co-sedimentation assay. In the presence of hSPACA9, we observed a clear increase in polymerized tubulin in the pellet fraction (Fig. S4B), indicating enhanced microtubule assembly. This effect was especially pronounced at lower tubulin concentrations, suggesting that hSPACA9 reduces the critical concentration for microtubule polymerization by facilitating nucleation and stabilizing early assembly intermediates.

### Microtubule binding and activity of hSPACA9 are mediated by its C-terminal tail

Next, we sought to identify the determinants underlying the interaction between hSPACA9 and microtubules. hSPACA9 is composed of eight α-helices arranged into four α-helix bundle (H1-H4), with insertions following H1 and H3, and a disordered C-terminal tail extending beyond the structured core (Fig. 4A). The microtubule-binding surface of hSPACA9 is formed by the short helices (25). To gain insight into the interaction of hSPACA9 with the microtubules lattice, we generated a mutant in the conserved residues Q20A, Q21A, T61A, R64A and N123A, which were identified as key determinants of microtubule binding (25). The mutant, hSPACA9^5A^, was tested for its ability to interact with microtubules. Surprisingly, a co-sedimentation assay confirmed the interaction between hSPACA9^5A^ and microtubules (Fig. 4B). In contrast, a pull-down assay with soluble tubulin showed no detectable interaction, consistent with the behavior of hSPACA9 (Fig. S2). We further examined whether the hSPACA9^5A^ mutant altered microtubule dynamics; however, its effects were indistinguishable from those of the wild-type protein (Fig. 4C). Specifically, no significant changes were observed in growth rate (12.24 ± 0.77 nm/sec vs. 10.42 ± 0.81 nm/sec), shrinkage rate (66.3 ± 7.11 nm/sec vs. 59.28 ± 5.1 nm/sec), catastrophe frequency (0.08 ± 0.01 events/min 0.05 ± 0.02 events/min) or rescue frequency compared to wild-type hSPACA9 (3.5 ± 0.44 events/min vs. 2.9 ± 0.36 events/min) at similar concentration, respectively. Consistently, microtubule extensions in the presence of hSPACA9^5A^ were comparable to those with hSPACA9 (Fig. S5). Thus, under our assay conditions, these residues are not required for hSPACA9-microtubule interaction or for modulating microtubule dynamic instability.

**Figure 4.**
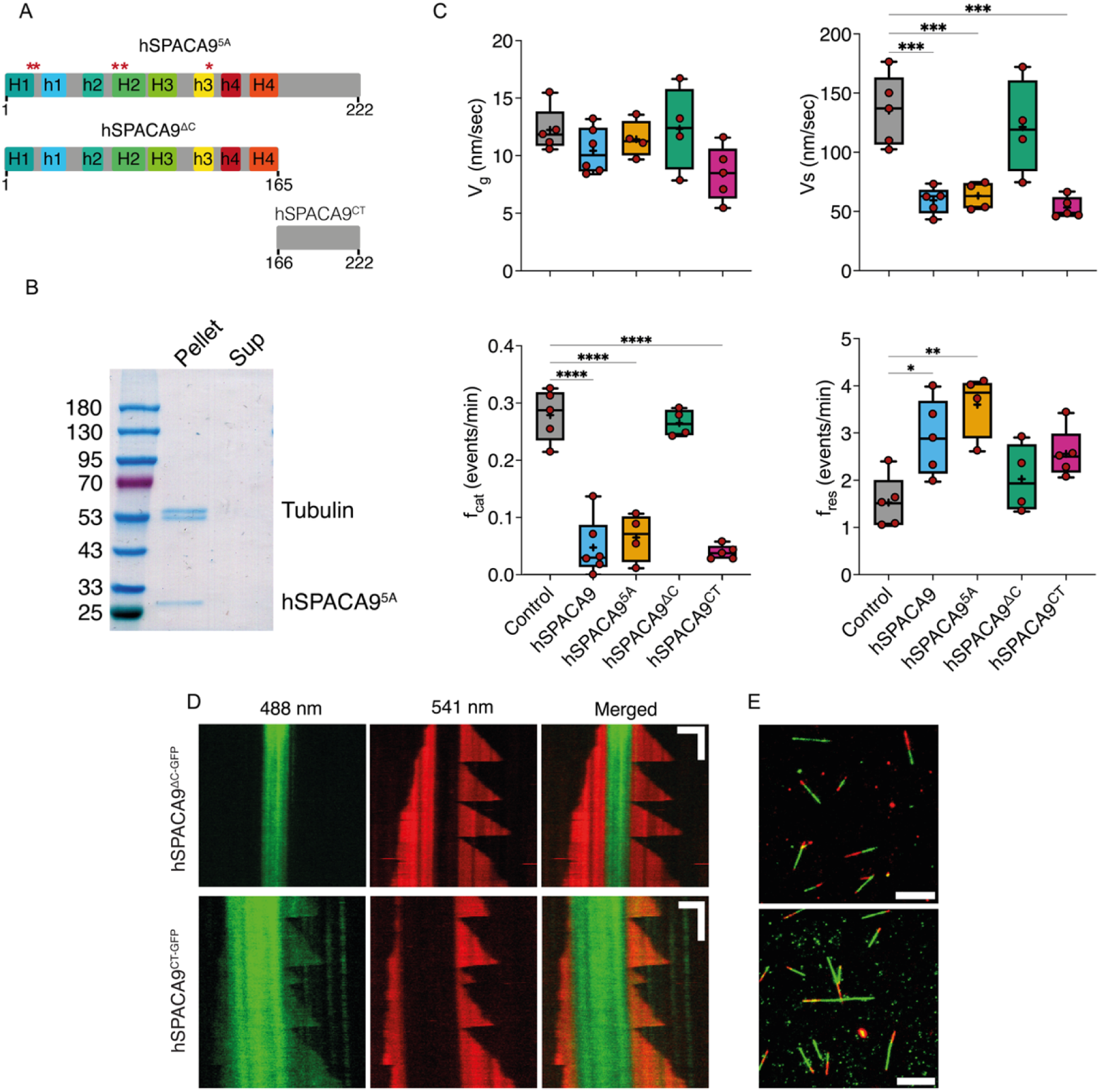
The C-terminal tail of hSPACA9 is essential for its binding to microtubules. (**A**) Schematic representation of full-length hSPACA9 and its truncated versions. Red asterisks indicate the locations of mutations. (**B**) Coomassie Blue-stained SDS-PAGE analysis of a co-sedimentation assay showing the binding of hSPACA9^5A^ to microtubules in the pellet. (**C**) Quantification of microtubule dynamic properties in the absence of hSPACA9 (control) and in the presence of hSPACA9, hSPACA9 with mutations (hSPACA9^5A^), hSPACA9 lacking the C-terminal region (hSPACA9^ΔC^), and C-terminal fragment (hSPACA9^CT^). (**D**) Representative kymographs of microtubules in the presence of 300nM hSPACA9^ΔC-GFP^, showing no binding of the truncated version to microtubules. In contrast, 300nM hSPACA9^CT-GFP^ binds microtubule. Scale bars: 5 µm, 3 min. (**E**) Representative TIRFM images of dynamic microtubules in the presence of 300nM hSPACA9^ΔC-GFP^ (top) and 300nM hSPACA9^CT-GFP^ (bottom) after 15 min. Scale bar: 5 µm. n =188, 226; data pooled across at least 4 independent experiments. *P< 0.05; **P< 0.01; ***P< 0.001 ;****P< 0.0001 as determined by one-way ANOVA followed by Dunnett’s multiple comparisons test.

If the short helices are not the primary determinant of hSPACA9-microtubule binding, what is? We addressed this question using truncation constructs. Initial attempts to generate N-terminally truncated versions of hSPACA9 resulted in constructs that were highly susceptible to degradation, likely due to disruption of the hydrophobic core of the helix bundle. We therefore focused on the C-terminal region and generated a shorter construct, hSPACA9^ΔC^, lacking the disordered tail (Fig. S6A). Surprisingly, no significant changes in microtubule dynamic properties compared to the control without hSPACA9 were observed, as reflected in the growth rate (12.33 ± 1.82 nm/sec), shrinkage rate (121.3 ± 20.17 nm/sec) catastrophe frequency (0.26 ± 0.01 events/min) or rescue frequency (2.02 ± 0.36 events/min) (Fig. 4C, movie S7). To further validate these results, we expressed and purified hSPACA9^ΔC-GFP^ (Fig. S6B) and found that it failed to interact with microtubules (Fig. 4D, movie S8). To determine if the C-terminus of hSPACA9 interacts directly with the microtubule lattice, we expressed and purified the C-terminus of hSPACA9 (166–222) either fused to GFP (hSPACA9^CT-GFP^) or untagged (hSPACA9^CT^) (Fig. S6C and D). TIRFM imaging revealed that hSPACA9^CT-GFP^ interacts directly with the microtubule lattice, but this interaction has no effect on microtubule dynamics (Fig. S7, movie S9). A pelleting assay confirmed that the unlabeled hSPACA9^CT^ construct also binds microtubules (Fig. S8). However, unlike hSPACA9^CT-GFP^, hSPACA9^CT^ altered microtubule dynamic instability, exhibiting effects comparable to those observed for full-length hSPACA9, with growth rate (8.45 ± 1.05 nm/sec), shrinkage rate (53.3 ± 3.92 nm/sec) catastrophe frequency (0.04 ± 0.01 events/min) and rescue frequency (2.56 ± 0.23 events/min) (Fig 4C – E, movie S10). Thus, the C-terminal region is sufficient to mediate microtubule binding and to recapitulate the effects of hSPACA9 on microtubule dynamic instability *in vitro*.

### hSPACA9 modulates shrinkage but not lattice breakage

Our results show that hSPACA9 supports microtubule growth by inhibiting dynamic instability, raising the question of whether it also provides protection against mechanical forces generated by the dynein arms during ciliary beating or by the IFT machinery. Indeed, both *in vitro* and *in vivo* studies have recently demonstrated that motor proteins induce damage to the microtubule lattice (36–38). To understand whether hSPACA9 stabilizes the microtubule lattice against mechanical stress induced by motor proteins, we performed motility assays on capped GDP-microtubules (Fig. 5A and B) (36).

**Figure 5.**
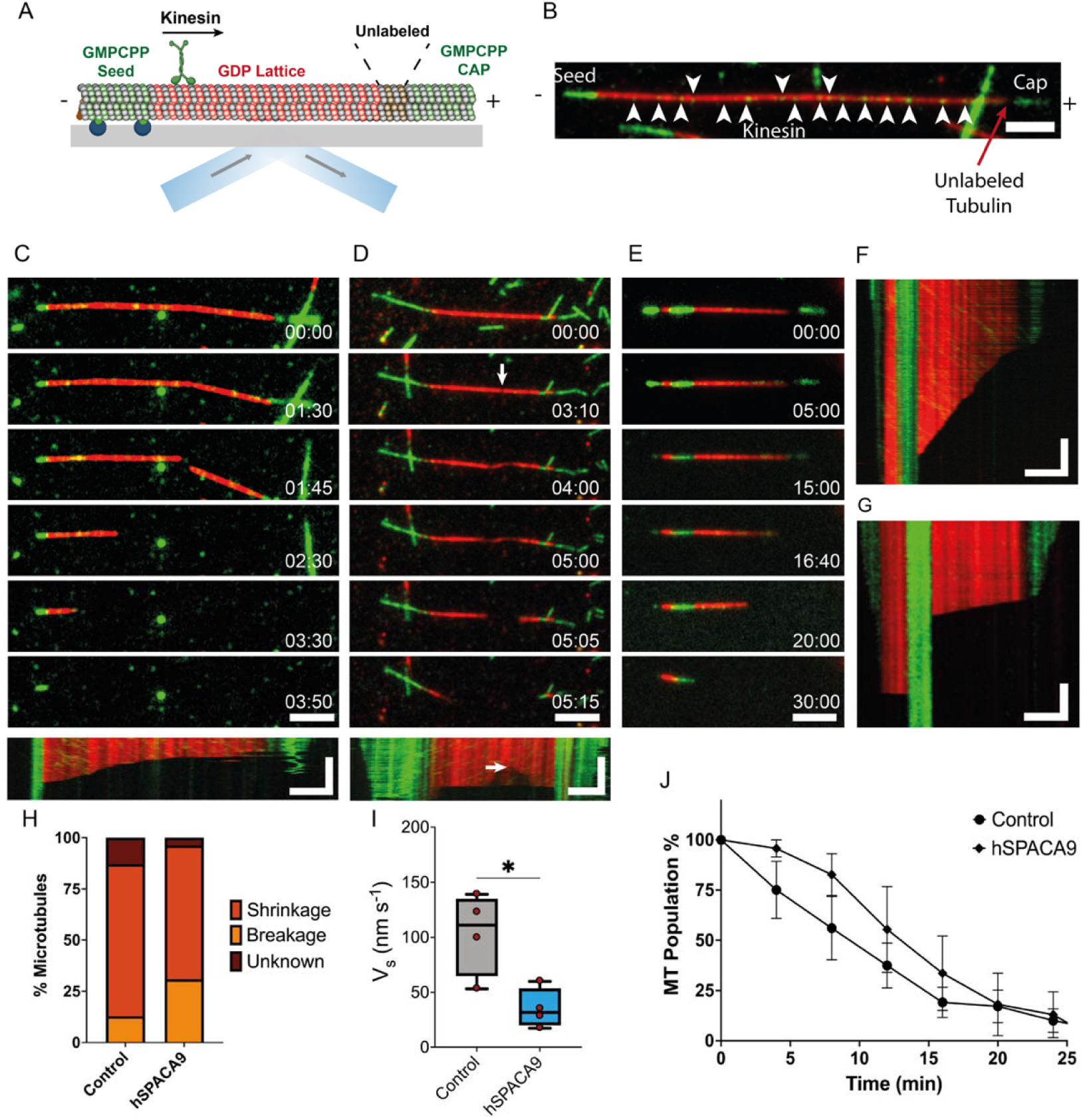
hSPACA9 has no effect on microtubule stability after motor protein-induced damage. (**A**) Schematic illustrating the TIRFM-based motility assay. A GMPCPP-stabilized seed (green) is anchored to the coverslip via an avidin-biotin interaction. After GDP-microtubules (red) polymerize from the seed, free TAMRA-labeled tubulin is washed away using unlabeled tubulin (brown) before capping with GMPCPP-tubulin (green). (**B**) TIRFM image of a microtubule in the motility assay. Scale bar: 3 μm (**C**) Time-lapse series and kymograph showing the sudden breakage of a GDP-microtubule in the presence of 10 nM GFP-tagged kinesin-1. (**D**) Time-lapse series and kymograph showing the progressive breakage of a GDP-microtubule in the presence of 10 nM GFP-tagged kinesin-1. (**E**) Time-lapse series showing the depolymerization of the cap and GDP-microtubule in the presence of 10 nM GFP-tagged kinesin-1. All scale bars: 3 μm, 3 min (**F**) Representative kymographs illustrating microtubule depolymerization in the presence of hSPACA9 (**G**) Representative kymographs illustrating microtubule depolymerization in the absence of hSPACA9. Scale bars: 3 μm, 3 min (**H**) Bar plot showing the percentage of microtubules exhibiting shrinkage, breakage, or undetermined events. Data pooled from 4 independent experiments. (**I**) Box plot depicting the shrinkage rate in the absence (control) and presence of hSPACA9. (**J**) Percentage of surviving microtubule population. n = 107, 109; data pooled from 4 independent experiments.

In both the presence and absence of hSPACA9, we observed damage leading to breakage within the GDP-microtubule shaft (Fig. 5C – D, movies S11-S13), as well as microtubule shrinkage (Fig. 5E, movie S14). Notably, hSPACA9 occasionally induced a slow expansion of the damaged region over several minutes, ultimately resulting in the microtubule splitting into two distinct fragments (Fig. 5D, movie S13). This phenomenon was not observed in the absence of hSPACA9. In all conditions, breakage was restricted to the GDP lattice; no breakage events were detected at the GMPCPP-stabilized seed or cap.

Most microtubule damage originated as shrinkage from the capped microtubule end, with over 75% of events initiating at the plus end, regardless of cap size (Fig. 5E – H, movie S14). In contrast, no significant shrinkage was observed from the stabilized seed, consistent with previous findings that the binding of the seed to the surface protects the microtubule from damage (39). The shrinkage rate in the presence of hSPACA9 was significantly slower (37.53 ± 4.09 nm/sec) compared to its absence (91.88 ± 7.61 nm/sec) (Fig. 5I), and comparable to our dynamic instability assays that included free tubulin (Fig. 2B). Despite its ability to slow shrinkage, hSPACA9 had no significant effect on overall microtubule survival (Fig. 5J). Thus, hSPACA9 appears to modulate lattice mechanics by delaying shrinkage and influencing damage propagation, rather than acting as a direct barrier to microtubule breakage.

## Discussion

MIPs constitute a distinct class of ciliary proteins conserved across eukaryotic species (10). This conservation underscores their fundamental importance; however, the striking structural and compositional diversity among MIPs suggests that they have evolved specialized roles tailored to the specific functional demands of different cilia types. In this study, we investigated the molecular basis underlying the role of hSPACA9, a MIP conserved among vertebrates, in regulating microtubule dynamics. Our results reveal that hSPACA9 contains two functional regions: an unstructured C-terminal tail that mediates microtubule binding and modulates microtubule dynamic instability, and a structured core that likely contributes to the architectural organization of the protein within the microtubule lumen. Although previous cryo-EM reconstructions positioned the core domain of hSPACA9 along the microtubule lumen, the C-terminal tail was not resolved in these structures and the binding activity of individual domains was not assessed (25, 28). In contrast, our TIRFM experiments show that the C-terminal tail is both necessary and sufficient for microtubule binding, whereas the core domain alone does not interact with the lattice. The tail is predicted to be intrinsically disordered based on its amino acid composition, which is enriched in disorder-promoting residues such as proline and glycine. Sequence analysis of multiple SPACA9 orthologs revealed several short, conserved motifs within the disordered tail, suggesting that this region may contribute to microtubule interaction (Fig. S9). Thus, the tail may act as a flexible tether that promotes the retention of hSPACA9 within the microtubule lumen. The presence of conserved residues within a disordered domain further implies that this interaction may involve sequence-specific recognition, rather than nonspecific interaction.

Furthermore, while conserved residues in the core domain have been proposed to mediate microtubule interactions (25), our findings indicate that these residues do not affect either microtubule binding or dynamic properties (Fig. 4C). Despite their evolutionary conservation and apparent structural relevance, simultaneous mutation of all five residues did not impair hSPACA9 binding to microtubules or its regulatory activity. Several possibilities may explain this discrepancy: (1) the interaction interface may be more distributed or functionally redundant than previously appreciated, such that disrupting a subset of residues does not abolish binding; (2) the mutations may not have substantially perturbed the local structure or charge landscape; or (3) binding may be predominantly mediated by the disordered C-terminal tail under our assay conditions, effectively masking the contribution of the structured domain. Rather than conflicting with the structural model, these results suggest that the observed interactions may be permissive or stabilizing rather than essential, and that the tail may play a primary role in initial microtubule engagement. Further mutational and functional dissection will be required to define the minimal elements necessary for hSPACA9 binding and activity.

hSPACA9 has been shown to bind two adjacent protofilaments, suggesting a potential role in stabilizing lateral interactions within the microtubule lattice (25). Our findings support this notion, demonstrating that hSPACA9 reduces catastrophe events and increases rescue frequency (Fig. 2). Furthermore, the reduced shrinkage rate likely contributes to microtubule stability by extending the window for rescue and facilitating recovery following catastrophes. In contrast, hSPACA9 has no significant effect on the microtubule growth rate, which can be explained by its inability to bind free tubulin. Together, these results suggest that hSPACA9 functions as a stabilizing element, effectively acting as a molecular staple that holds protofilaments together. Furthermore, our TIRFM assays and TEM images indicate that hSPACA9 stabilizes protofilaments at the growing microtubule ends (Fig. 3), which may have implications for the insertion mechanisms of MIPs into the microtubule lumen and the formation of nascent microtubule formation (Fig. 6). Previously, it was shown that ciliary tubulin tends to form sheet-tip structures, and our findings suggest that hSPACA9 contributes to stabilizing these sheets, potentially facilitating their transition into a closed microtubule lattice (40).

**Figure 6.**
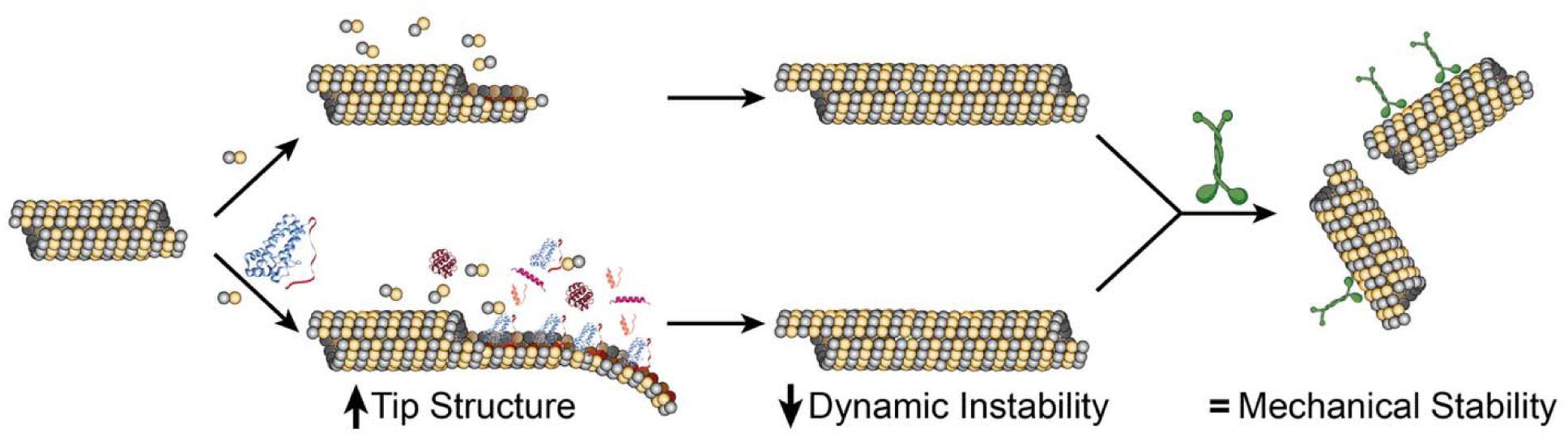
Suggested Model. In the absence of hSPACA9, microtubules exhibit dynamic instability, characterized by frequent growth and shrinkage events, and form only short tip structures until the full microtubule is assembled. Motor proteins induce damage to the microtubule shaft, potentially leading to breakage. In contrast, hSPACA9 supports microtubule growth, suppresses dynamic instability, and stabilizes protofilaments, resulting in an extended tip structure. However, hSPACA9 does not protect against mechanical stress generated by motor proteins. The structure of hSPACA9 was adapted from PDB ID: 7UNG.

Finally, our mechanical stability assays provide new insight into how hSPACA9 influences microtubule behavior under motor-induced stress. While similar assays have been used previously to demonstrate that motor proteins can damage the microtubule lattice, this study represents, to our knowledge, the first application of such an assay to assess the stabilizing effects of a microtubule-associated protein. Despite the lack of significant protection against breakage, hSPACA9 consistently reduced the rate of shrinkage following damage, suggesting a partial stabilizing effect on the GDP lattice (Fig. 6). It is also important to consider that hSPACA9 functions in the context of a dense network of microtubule inner proteins. Although our assays focused exclusively on hSPACA9, it is likely that multiple MIPs act cooperatively to reinforce axonemal microtubules under mechanical stress. In particular, filamentous MIPs (fMIPs), which extend across multiple protofilaments or span several tubulin subunits, may exert a more pronounced stabilizing effect by physically linking or constraining lattice segments.

Importantly, because hSPACA9 is relatively small and contains multiple lysine and cysteine residues positioned within the α-helical core and putative tubulin-interaction surfaces, common labeling strategies (e.g., NHS-ester or maleimide chemistry) risk steric or chemical perturbation and heterogeneous conjugation. To mitigate these limitations, we instead employed a C-terminal GFP fusion; however, even this strategy altered microtubule dynamics compared to the untagged protein (Fig. 1, Fig. S7). A similar phenomenon has been reported for other microtubule-associated proteins (41, 42). Given the periodic intralumenal organization of hSPACA9, we suspect that steric hindrance introduced by large tags, such as GFP, interfere with its interaction with the microtubule lattice. While our experiments specifically used GFP, this effect is likely relevant to other large tags. These observations underscore the need to carefully consider tag size and positioning when assessing MIP localization and function.

The stabilization mechanism of hSPACA9 expands the range of MIP-mediated microtubule regulation, highlighting the diversity of strategies used to reinforce microtubules. These findings highlight that different MIPs employ distinct strategies to achieve microtubule stabilization. For example, FAP85 reduces the depolymerization rate, contributing to enhanced stability (43). The DMT inner junction protein FAP20 not only increases growth rates, which can facilitate fast recovery from catastrophe events, but also reduces catastrophe frequency (44). In addition, both FAP20 and PACRG, which form the inner junction proteins, have been shown to recruit tubulin to the microtubule lattice, potentially contributing to the formation of the B-tubule (44, 45). Similarly, the ciliary tip protein CSPP1 stabilizes microtubules by binding to pre-catastrophe microtubule ends, preventing full catastrophes by suppressing shrinkage and inducing pauses until regrowth resumes (46). Furthermore, CSPP1 was found to enter the damaged microtubule lumen, where it supports microtubule lattice repair by stabilizing rescue sites and enabling tubulin incorporation.

Overall, our results highlight hSPACA9 as a distinct stabilizing element within the axonemal microtubule, contributing to both dynamic and mechanical regulation of the lattice. The functional diversity among MIPs, as exemplified by hSPACA9, suggests a modular organization in which different proteins support specific aspects of microtubule properties and architecture. Future work should explore how MIPs act in concert with other MIPs, including potential cooperative mechanisms between hSPACA9 and stabilizers of axonemal microtubules (SAXO) proteins that colocalize (25), to maintain microtubule integrity under mechanical stress. Such efforts will be essential to fully unravel the molecular logic of ciliary architecture and its implications for motility. Importantly, although this study focuses on a single MIP in a simplified system, it provides a mechanistic framework for understanding how individual MIPs contribute to the regulation of microtubule dynamics and, more broadly, to the organization of the axonemal lattice.

## Materials and Methods

### DNA constructs

The coding sequence for human SPACA9 (accession number NP_001303826.1) was purchased from Integrated DNA Technologies and inserted into the pET22 vector using Gibson assembly (NEB). The construct included a C-terminal GFP tag followed by a 6×His tag. Untagged hSPACA9 was generated by inserting the hSPACA9 sequence into the pET22 vector without the GFP tag, also using Gibson assembly. Mutants of hSPACA9-GFP, containing a C-terminal EGFP tag followed by a 6×His tag, were ordered from Genescript using the same pET22 vector.

### Protein Expression & Purification

All protein constructs were expressed in *E. coli* BL21 cells overnight at 16°C. Cell pellets were resuspended in lysis buffer (50 mM sodium phosphate pH 7.0, 250 mM NaCl, 1mM MgCl_2_, 0.1% Triton X-100, 1 mM DTT, 1 mg/mL lysozyme, 20 mM imidazole, 10uL Benzonase (Sigma-Aldrich) supplemented with EDTA-free protease inhibitors (Sigma-Aldrich)) and lysed in a glass dounce homogenizer. The crude lysate was clarified by centrifugation at 30,000×g for 40 minutes at 4°C. Protein purifications were performed using an AKTA go (Cytiva Life Sciences) at 4°C. The clarified lysate was applied to a HisTrap HP column (Cytiva Life Sciences) following the manufacturer’s protocol and eluted with 50 mM NaH_2_PO_4_ (pH 7.0), 250 mM NaCl, 1 mM DTT, 1 mM MgCl_2_ and 450 mM imidazole. The fractions were analyzed by SDS-PAGE and hSPACA9-rich Ni-NTA fractions were subsequently subjected to size-exclusion chromatography (SEC) using a Superdex 200 increase 10/300 column (Cytiva Life Sciences) pre-equilibrated in BRB80 (80 mM PIPES, pH 6.9, 1 mM EGTA, 1 mM MgCl_2_). All purified proteins were aliquoted and snap-frozen in liquid nitrogen and stored at −80°C.

### C-Terminal Tail Expression & Purification

Tobacco Etch Virus (TEV) protease recognition sequence was inserted into the coding region of hSPACA9, upstream the C-terminal tail. hSPACA9-TEV was expressed and purified using Ni-affinity chromatography. Following purification, the recombinant protein was subjected to TEV protease cleavage, yielding three species: the uncleaved full-length hSPACA9 and two cleavage products (hSPACA9^ΔC^ and hSPACA9^CT^). The reaction mixture was subsequently applied to a second Ni-affinity chromatography step to separate tagged components, followed by final purification by SEC using a Superdex 70 increase 10/300 column (Cytiva Life Sciences).

### Tubulin and Microtubule Preparation

GMPCPP-stabilized microtubules were prepared as described previously (33). A tubulin mix consisting of 70% unlabeled porcine brain tubulin, 20% biotin-labeled porcine tubulin, and 10% fluorescently-labeled porcine tubulin (Cytoskeleton, Inc.) was used. Fresh GMPCPP microtubules were prepared by incubating the tubulin mix with 1 mM GMPCPP (Jena Biosciences) at 37°C for 30 minutes. Polymerized microtubule seeds were pelleted by centrifugation at 21,300×g for 30 minutes at 35°C and resuspended in warm BRB80. Seeds were flash frozen in liquid nitrogen and stored at -80°C until use.

### Microtubule Cosedimentation Assay

Microtubules were polymerized by incubating 12 μM porcine tubulin with increasing amounts of hSPACA9 or hSPAC9-GFP in BRB80 buffer with 10 mM GTP at 37°C for 30 minutes. After polymerization, 25 μM taxol was added to stabilize the microtubules. The taxol-stabilized microtubules were then pelleted by centrifugation at 21,300×g for 30 minutes at 35°C and resuspended in SDS-PAGE sample buffer.

### SDS-PAGE and Immunoblot

Samples were boiled for 5 min in SDS–PAGE sample buffer and resolved on an SDS-PAGE gels that were stained in Coomassie blue. The purified C-terminal tail fragment, owing to its low molecular weight, was resolved using 16.5% Tricine-SDS-PAGE to ensure optimal separation. For Western blot analysis, the following primary antibodies were used: anti-acetylated α-tubulin (clone 6–11B-1; 1:50,000; MilliporeSigma); anti α-tubulin (clone B-5-1-2; 1:1000; MilliporeSigma).

### TIRF microscopy

Imaging was performed using a Nikon Eclipse Ti2-E microscope with a perfect focus system (Nikon). The system was equipped with a Prime BSI sCMOS camera (Teledyne Photometrics) and a LightHUB+ laser engine (488/561/638 nm; Omicron), utilizing a 100×/1.49 N.A. CFI Apo TIRF oil objective. Image acquisition was controlled by NIS-Elements software (Nikon). Microscope chambers were constructed as previously described (32). Briefly, 22×22 mm and 18×18 mm coverslips were cleaned using piranha solution and silanized. Coverslips were separated by Parafilm strips to create a flow channels for the exchange of solution. For dynamic instability assays, seeds were attached to silanized glass slides via biotin-neutravidin interactions. Imaging buffer containing concentrations of 10 µM tubulin, and proteins at the concentrations indicated in the figure legend were introduced into the imaging chamber. Imaging buffer supplemented with oxygen scavengers (BRB80, 80 mM glucose, 8 μg/ml glucose oxidase, 32 μg/ml catalase, 20 µg/ml casein, 20 µM dithiothreitol and 1 mM GTP). Dynamic assays were performed at 30°C and imaged at frame rate of 0.2 fps. Images were acquired using NIS-Elements (Nikon).

### Transmission Electron Microscopy

For negative-stain TEM, 5 μL of sample was applied to glow-discharged formvar/carbon-coated copper grids (Electron Microscopy Sciences) and allowed to adsorb for 1 min. Excess sample was blotted, and grids were stained with 2% (w/v) uranyl acetate for 2 min. The stain was blotted off, and grids were washed three times with distilled water prior to air-drying. Micrographs were acquired using a JEOL JEM-120i transmission electron microscope operated at 120 kV.

### Subtilisin Treatment of Microtubules

Polymerized microtubules were treated with subtilisin to remove the C-terminal tails (E-hooks) of tubulin as described previously. Briefly, assembled microtubules were incubated with subtilisin (Sigma-Aldrich) at a final concentration of 200 μg/mL for 20 min at 30 °C. The reaction was terminated by addition of 4-(2-Aminoethyl)benzenesulfonyl fluoride hydrochloride (Thermo Scientific) to a final concentration of 20 mM. Treated microtubules were immediately used after centrifugation at 21,300× g for 30 min to remove residual protease.

### Circular Dichroism (CD) spectroscopy

CD spectra were recorded at 20 °C using a Chirascan spectropolarimeter (Applied Photophysics). Measurements for hSPACA9 and hSPACA9^ΔC^ were performed at a protein concentration of 15 μM in phosphate buffer using a 0.1-cm path-length quartz cuvette. Far-UV spectra were collected from 190 to 260 nm with a step size of 1 nm and a bandwidth of 1 nm.

### Microtubule tip curvature visualization

To compare the curvature of microtubule tips, individual tip contours were manually traced in Fiji and exported as XY coordinate profiles using the Kappa plugin for visualization. Each trace was aligned so that the initial point was positioned at the origin (0,0), and the curves were uniformly rotated to orient their elongation axis along the negative X-direction, standardizing both directionality and curvature across all samples. To enhance visual clarity without altering structural information, curves were evenly distributed around the origin using additional display rotations. All curves were plotted using a custom MATLAB script with a colorblind-friendly palette.

### Motility assay

For motility assays, HiLyte 488 microtubule seeds stabilized with GMPCPP and containing 10% biotinylated-tubulin were attached to silanized glass slides via biotin-neutravidin interactions. Unbound microtubules were washed with warm BRB80, and GDP-microtubules were polymerized for 5 minutes in BRB80 supplemented with 14 µM TAMRA-labeled tubulin and 1 mM GTP. Free TAMRA-labeled tubulin was then washed away using warm BRB80 supplemented with 10 µM unlabeled tubulin, and immediately the microtubules were capped for 1 minute with 10 µM HiLyte 488 tubulin in the presence of 1 mM GMPCPP and 2 mM of MgCl_2_. Finally, free tubulin was washed away with warm BRB80, and 10 nM kinesin-1 was added in the presence of 1 mM ATP in oxygen scavenger solution. Motility assays were performed at 30°C and imaged at frame rate of 0.2 fps. Images were acquired using NIS-Elements (Nikon).

### Pull-Down Assay

Triple-washed Ni Sepharose 6 Fast Flow beads were equilibrated in binding buffer (50 mM NaH_2_PO4, 300 mM NaCl, 10 mM imidazole, pH 7.0). Protein was then added and incubated with gentle rotation overnight at 4°C. Beads were pelleted by centrifugation and washed three times with 500 µl binding buffer. Bound proteins were eluted with 300 mM imidazole, and samples were analyzed by SDS-PAGE.

### Image analysis

Image analysis was performed by generating kymographs of individual MT seeds using ImageJ analysis software (National Institutes of Health) as described previously

(40). Kymographs were analyzed manually by fitting lines to kymographs of growing microtubules. Catastrophe frequency was calculated by dividing the number of catastrophes by the total growth time. Rescue frequency was calculated by dividing the number of rescues by the total shrinkage time.

### Conservation Analysis

Evolutionary conservation of full-length hSPACA9 was analyzed using the ConSurf server (https://consurf.tau.ac.il). Homologues were identified with HMMER against the UniRef90 database (E-value cutoff 0.0001), and filtered for 35–95% sequence identity and ≥60% coverage. From 358 unique sequences, 150 were selected for analysis. Multiple sequence alignment was performed using MAFFT. Conservation scores were calculated using a Bayesian approach with the JTT substitution model. Scores were mapped along the hSPACA9 sequence to identify conserved regions.

### Statistical analysis

Data plotting and curve fitting were performed using Prism 10 (GraphPad). All quantification and statistical analyses, including the statistical tests used and the exact values of N (number of trials) and n (number of microtubules), are detailed in the respective figure legends. All experiments were conducted at least three times. Differences were considered significant at *p < 0.05, **p < 0.01, ***p < 0.001, and ****p < 0.0001.

## Acknowledgments

We thank Lior Giterman and Helena Sabanay for their assistance with TEM imaging, and Dr. Yulia Shenberger for support with CD measurements. We are grateful to Prof. Moshe Dessau for his valuable advice and insightful discussions, and to Albert Mostovoy for technical support. We also thank Prof. Luke Rice, Prof. Leah Gheber, Dr. Iris Grossman-Haham, and all lab members for their critical reading of the manuscript. This work was supported in part by the Israel Science Foundation (ISF) under grants 2251/23 and 2741/23 (R.O.).

## Author contributions

Conceptualization R.O.; investigation, M.A., S.F.B-U; funding acquisition R.O; supervision, R.O; writing R.O.

## References

1. D. R. Mitchell, Evolution of cilia. Cold Spring Harb. Perspect. Biol. 9, a028290 (2017).

2. G. B. Witman, K. Carlson, J. L. Rosenbaum, Chlamydomonas flagella. II. The distribution of tubulins 1 and 2 in the outer doublet microtubules. J. Cell Biol. 54, 540–555 (1972).

3. M. E. Porter, W. S. Sale, The 9 + 2 axoneme anchors multiple inner arm dyneins and a network of kinases and phosphatases that control motility. J. Cell Biol. 151, F37–F42 (2000).

4. D. Nicastro, et al., The molecular architecture of axonemes revealed by cryoelectron tomography. Science 313, 944–948 (2006).

5. T. Ishikawa, Axoneme structure from motile cilia. Cold Spring Harb. Perspect. Biol. 9, a028076 (2017).

6. D. Nicastro, et al., Cryo-electron tomography reveals conserved features of doublet microtubules in flagella. Proc. National Acad. Sci. U.S.A. 108, E845–E853 (2011).

7. N. Klena, G. Pigino, Structural Biology of Cilia and Intraflagellar Transport. Annu. Rev. Cell Dev. Biol. 38, 103–123 (2022).

8. I. Grossman-Haham, Towards an atomic model of a beating ciliary axoneme. Curr. Opin. Struct. Biol. 78, 102516 (2023).

9. M. Ichikawa, K. H. Bui, Microtubule Inner Proteins: A Meshwork of Luminal Proteins Stabilizing the Doublet Microtubule. BioEssays 40, 1700209 (2018).

10. M. Gui, R. Orbach, Microtubule inner proteins – bridging structure and function in ciliary biology. J. Cell Sci. 138 (2025).

11. S. E. Lacey, G. Pigino, The intraflagellar transport cycle. Nat. Rev. Mol. Cell Biol. 1–18 (2024). 10.1038/s41580-024-00797-x.

12. V. VanBuren, L. Cassimeris, D. J. Odde, Mechanochemical model of microtubule structure and self-assembly kinetics. Biophys. J. 89, 2911–2926 (2005).

13. L. M. Rice, E. A. Montabana, D. A. Agard, The lattice as allosteric effector: Structural studies of alpha beta- and gamma-tubulin clarify the role of GTP in microtubule assembly. Proc. National Acad. Sci. U.S.A. 105, 5378–5383 (2008).

14. G. J. Brouhard, L. M. Rice, Microtubule dynamics: an interplay of biochemistry and mechanics. Nat. Rev. Mol. Cell Bio. 19, 451–463 (2018).

15. T. Mitchison, M. Kirschner, Dynamic instability of microtubule growth. Nature 312, 237–242 (1984).

16. H. Bowne-Anderson, M. Zanic, M. Kauer, J. Howard, Microtubule dynamic instability: a new model with coupled GTP hydrolysis and multistep catastrophe. BioEssays 35, 452–461 (2013).

17. M. Ichikawa, et al., Subnanometre-resolution structure of the doublet microtubule reveals new classes of microtubule-associated proteins. Nat. Commun. 8, 15035 (2017).

18. M. Ma, et al., Structure of the decorated ciliary doublet microtubule. Cell 179, 909–922 (2019).

19. S. Kubo, et al., Native doublet microtubules from Tetrahymena thermophila reveal the importance of outer junction proteins. Nat. Commun. 14, 2168 (2023).

20. D. Stoddard, et al., Tetrahymena RIB72A and RIB72B are microtubule inner proteins in the ciliary doublet microtubules. Mol. Biol. Cell 29, 2566–2577 (2018).

21. A. A. Z. Khalifa, et al., The inner junction complex of the cilia is an interaction hub that involves tubulin post-translational modifications. ELife 9, e52760 (2020).

22. M. Gui, et al., De novo identification of mammalian ciliary motility proteins using cryo-EM. Cell 184, 5791–5806.e19 (2021).

23. M. M. Shimogawa, et al., FAP106 is an interaction hub for assembling microtubule inner proteins at the cilium inner junction. Nat. Commun. 14, 5225 (2023).

24. H. Lu, et al., Localisation and function of key axonemal microtubule inner proteins and dynein docking complex members reveal extensive diversity among vertebrate motile cilia. Development 151 (2024).

25. M. Gui, et al., SPACA9 is a lumenal protein of human ciliary singlet and doublet microtubules. Proc. Natl. Acad. Sci. U.S.A 119, e2207605119 (2022).

26. L. Tai, G. Yin, X. Huang, F. Sun, Y. Zhu, In-cell structural insight into the stability of sperm microtubule doublet. Cell Discov. 9, 116 (2023).

27. L. Zhou, et al., Structures of sperm flagellar doublet microtubules expand the genetic spectrum of male infertility. Cell 186, 2897–2910.e19 (2023).

28. M. R. Leung, et al., Structural specializations of the sperm tail. Cell 186, 2880–2896.e17 (2023).

29. Z. Chen, et al., De novo protein identification in mammalian sperm using in situ cryoelectron tomography and AlphaFold2 docking. Cell 186, 5041–5053.e19 (2023).

30. D. Zabeo, J. T. Croft, J. L. Höög, Axonemal doublet microtubules can split into two complete singlets in human sperm flagellum tips. FEBS Letters 593, 892–902 (2019).

31. M. R. Leung, et al., The multi-scale architecture of mammalian sperm flagella and implications for ciliary motility. EMBO J. 40, EMBJ2020107410 (2021).

32. C. Gell, et al., Microtubule dynamics reconstituted in vitro and imaged by single-molecule fluorescence microscopy. Methods Cell Biol. 95, 221–245 (2010).

33. W. G. Hirst, C. Kiefer, M. K. Abdosamadi, E. Schäffer, S. Reber, In Vitro Reconstitution and Imaging of Microtubule Dynamics by Fluorescence and Label-free Microscopy. Star Protoc. 1, 100177 (2020).

34. D. Chrétien, S. D. Fuller, E. Karsenti, Structure of growing microtubule ends: two-dimensional sheets close into tubes at variable rates. J. Cell Biol. 129, 1311–1328 (1995).

35. J. R. McIntosh, et al., Microtubules grow by the addition of bent guanosine triphosphate tubulin to the tips of curved protofilaments. J. Cell Biol. 265, jcb.201802138 (2018).

36. S. Triclin, et al., Self-repair protects microtubules from destruction by molecular motors. Nat. Mater. 20, 883–891 (2021).

37. B. G. Budaitis, et al., A kinesin-1 variant reveals motor-induced microtubule damage in cells. Curr. Biol. 32, 2416–2429.e6 (2022).

38. M. Andreu-Carbó, S. Fernandes, M.-C. Velluz, K. Kruse, C. Aumeier, Motor usage imprints microtubule stability along the shaft. Dev. Cell 57, 5–18.e8 (2022).

39. C. Tropini, E. A. Roth, M. Zanic, M. K. Gardner, J. Howard, Islands containing slowly hydrolyzable GTP analogs promote microtubule rescues. PLoS ONE 7, e30103 (2012).

40. R. Orbach, J. Howard, The dynamic and structural properties of axonemal tubulins support the high length stability of cilia. Nat. Commun. 10, 1838 (2019).

41. S. B. Skube, J. M. Chaverri, H. V. Goodson, Effect of GFP tags on the localization of EB1 and EB1 fragments in vivo. Cytoskeleton 67, 1–12 (2010).

42. J. M. Schrøder, et al., EB1 and EB3 promote cilia biogenesis by several centrosome-related mechanisms. J. Cell Sci. 124, 2539–2551 (2011).

43. J. Kirima, K. Oiwa, Flagellar-associated protein FAP85 is a microtubule inner protein that stabilizes microtubules. Cell Struct. Funct. 43, 1–14 (2018).

44. M. Bangera, A. Dungdung, S. Prabhu, M. Sirajuddin, Doublet microtubule inner junction protein FAP20 recruits tubulin to the microtubule lattice. Structure 31, 1535–1544.e4 (2023).

45. N. Khan, et al., Crystal structure of human PACRG in complex with MEIG1 reveals roles in axoneme formation and tubulin binding. Structure 29, 572–586.e6 (2021).

46. C. M. van den Berg, et al., CSPP1 stabilizes growing microtubule ends and damaged lattices from the luminal side. J. Cell Biol. 222, e202208062 (2023).

