## Supplementary material for "SPACA9 Acts as a Molecular Staple Modulating Microtubule Dynamic Instability": SI

### Supporting Information

**This PDF file includes:**

Figures S1 to S9

**Other supporting materials for this manuscript include the following:**

Movie S1. Microtubule dynamics in the absence of hSPACA9
Movie S2. Microtubule dynamics in the presence of hSPACA9
Movie S3. Microtubule dynamics in the presence of hSPACA9$^{GFP}$
Movie S4. Curved protofilament extensions at the microtubule tip
Movie S5. Rapid protofilament extension along dim microtubule tips in the presence of hSPACA9
Movie S6. Microtubule nucleation in the presence of hSPACA9
Movie S7. Microtubule dynamics in the presence of hSPACA9$^{\Delta C}$
Movie S8. Microtubule dynamics in the presence of hSPACA9$^{\Delta C\text{-}GFP}$
Movie S9. Microtubule dynamics in the presence of hSPACA9$^{CT\text{-}GFP}$
Movie S10. Microtubule dynamics with hSPACA9$^{CT}$ recapitulating full-length activity
Movie S11. Microtubule breakage during kinesin-driven motility in the absence of hSPACA9
Movie S12. Microtubule breakage during kinesin-driven motility in the presence of hSPACA9
Movie S13. Progressive microtubule breakage during kinesin-driven motility in the presence of hSPACA9
Movie S14. Microtubule shrinkage during kinesin-driven motility in the presence of hSPACA9

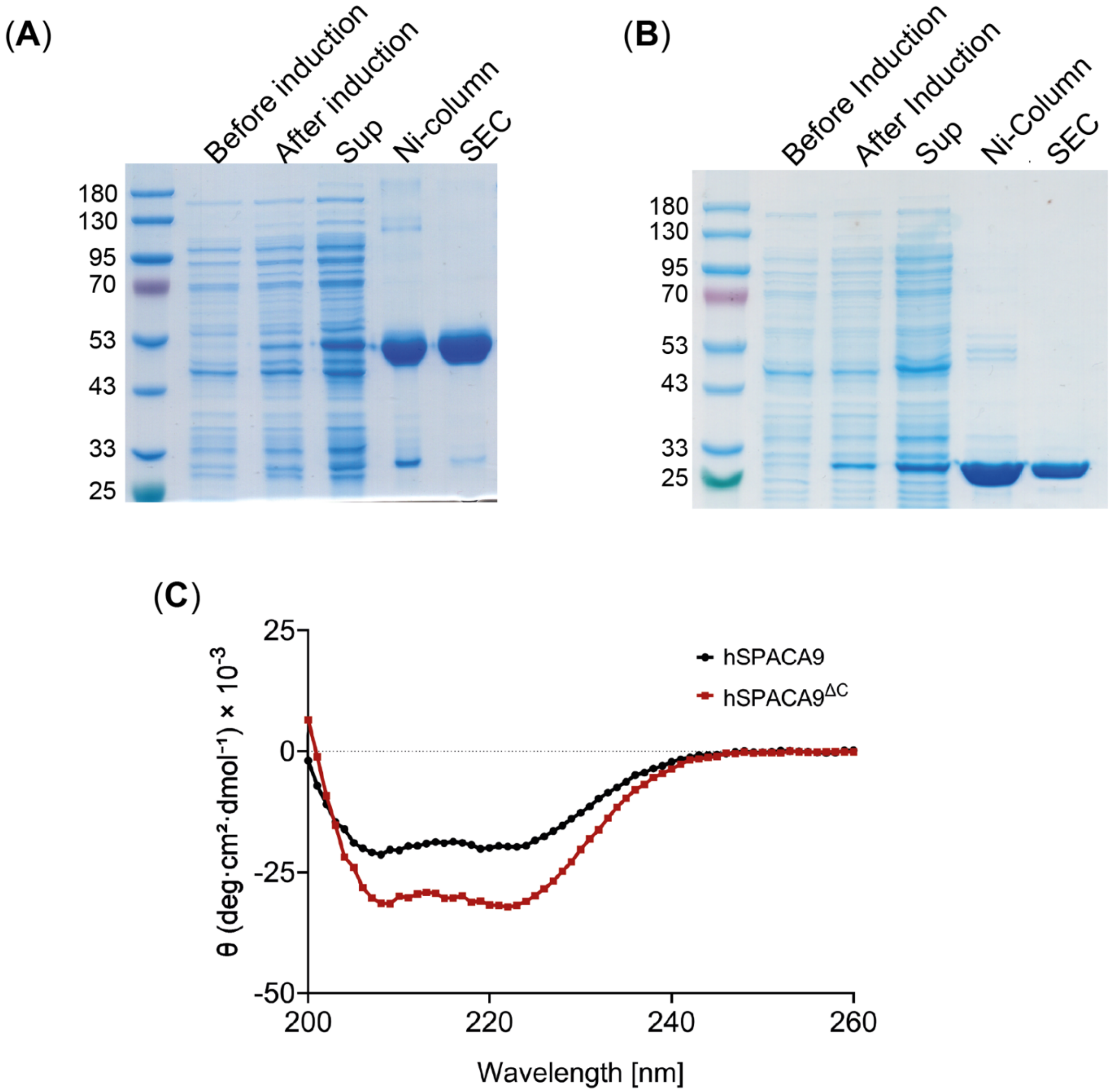


**Fig. S1. hSPACA9 purification and CD analysis**. SDS-PAGE gel, stained with Coomassie blue, showing the expression and purification steps for (**A**) hSPACA9$^{GFP}$ and (**B**) hSPACA9. Lanes from left to right: cell lysate before IPTG induction, cell lysate after IPTG induction, clarified supernatant (Sup), Ni-column elution, and size exclusion chromatography (SEC) elution. (**C**) Far-UV CD spectra of full-length hSPACA9 (black) and hSPACA9$^{\Delta C}$ (residues 1–165; red) recorded from 190–260 nm. Spectra exhibit characteristic minima at ~208 and ~222 nm indicative of α-helical secondary structure. Data below 200 nm were excluded from analysis due to increased buffer absorbance.

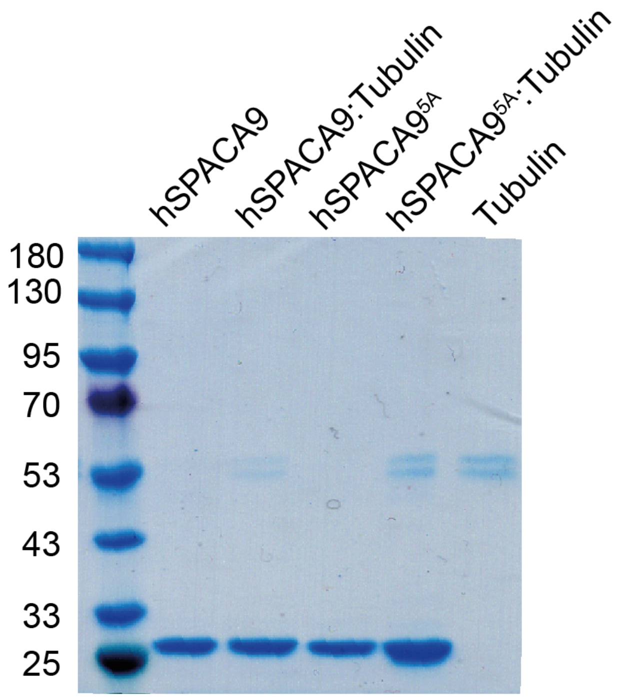


**Fig. S2**. **hSPACA9 does not bind tubulin in a pull-down assay**. Ni-NTA pull-down assay using His-tagged hSPACA9 to test for interaction with soluble tubulin. SDS-PAGE gel stained with Coomassie blue shows the pellet fractions from the following conditions: hSPACA9 alone, hSPACA9 with tubulin, hSPACA9$^{5A}$, hSPACA9$^{5A}$ with tubulin, and tubulin alone.

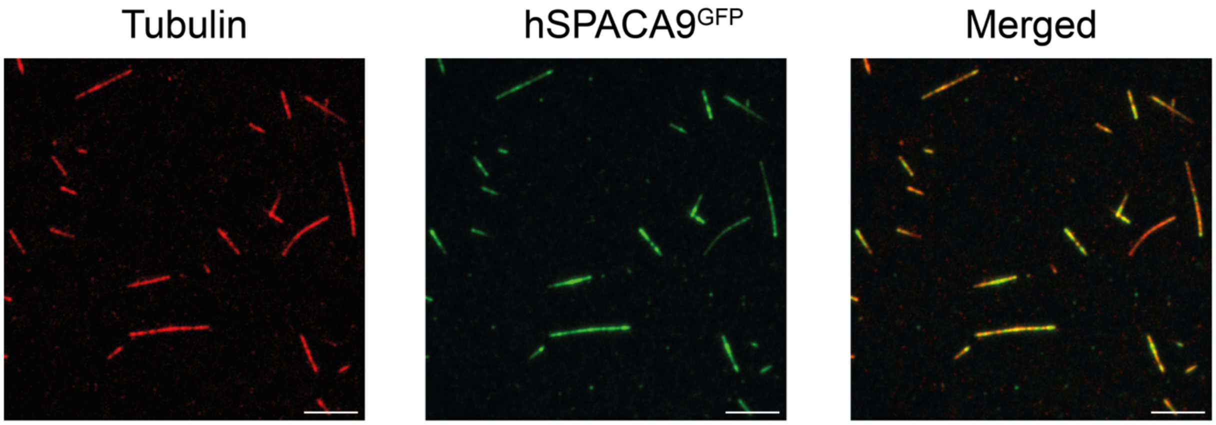


**Fig. S3. Subtilisin treatment of microtubules.** Microtubules were treated with subtilisin to proteolytically remove the C-terminal tails (E-hooks) of tubulin. Following treatment, hSPACA9$^{GFP}$ remained associated with the microtubules.

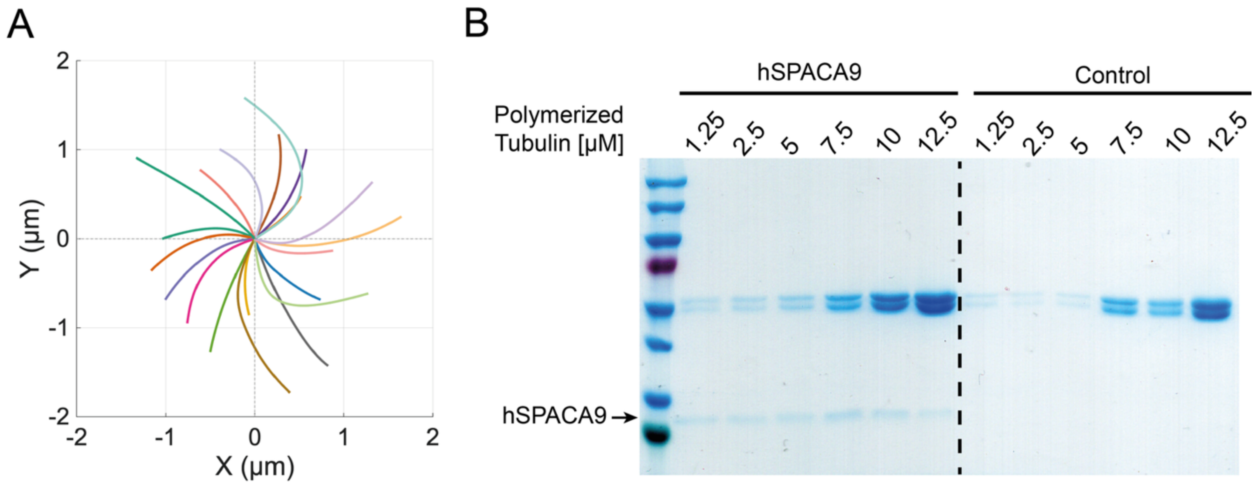


**Fig S4. Effect of hSPACA9 on microtubule tip structure and lattice binding. (A)** Graph illustrating individual traces of curved microtubule tip structures in the presence of 300nM hSPACA9. Each trace is color-coded to distinguish individual measurements. **(B)** Co-sedimentation (pelleting) assay of microtubules with hSPACA9. Increasing concentrations of polymerized tubulin were tested in the presence of 1 μM hSPACA9. Control samples under identical conditions lacked hSPACA9. In both cases, the samples correspond to the pellet fraction.

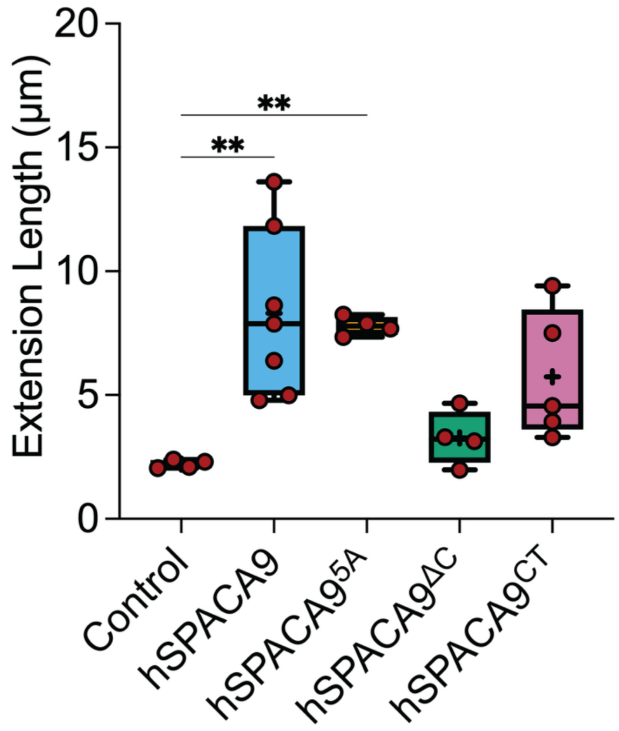


**Fig. S5. Effect of hSPACA9 constructs on microtubule extension length.** A plot showing the mean extension length after 15 min. Each dot represents the mean extension length from an individual experiment, while the plus sign indicates the overall mean.

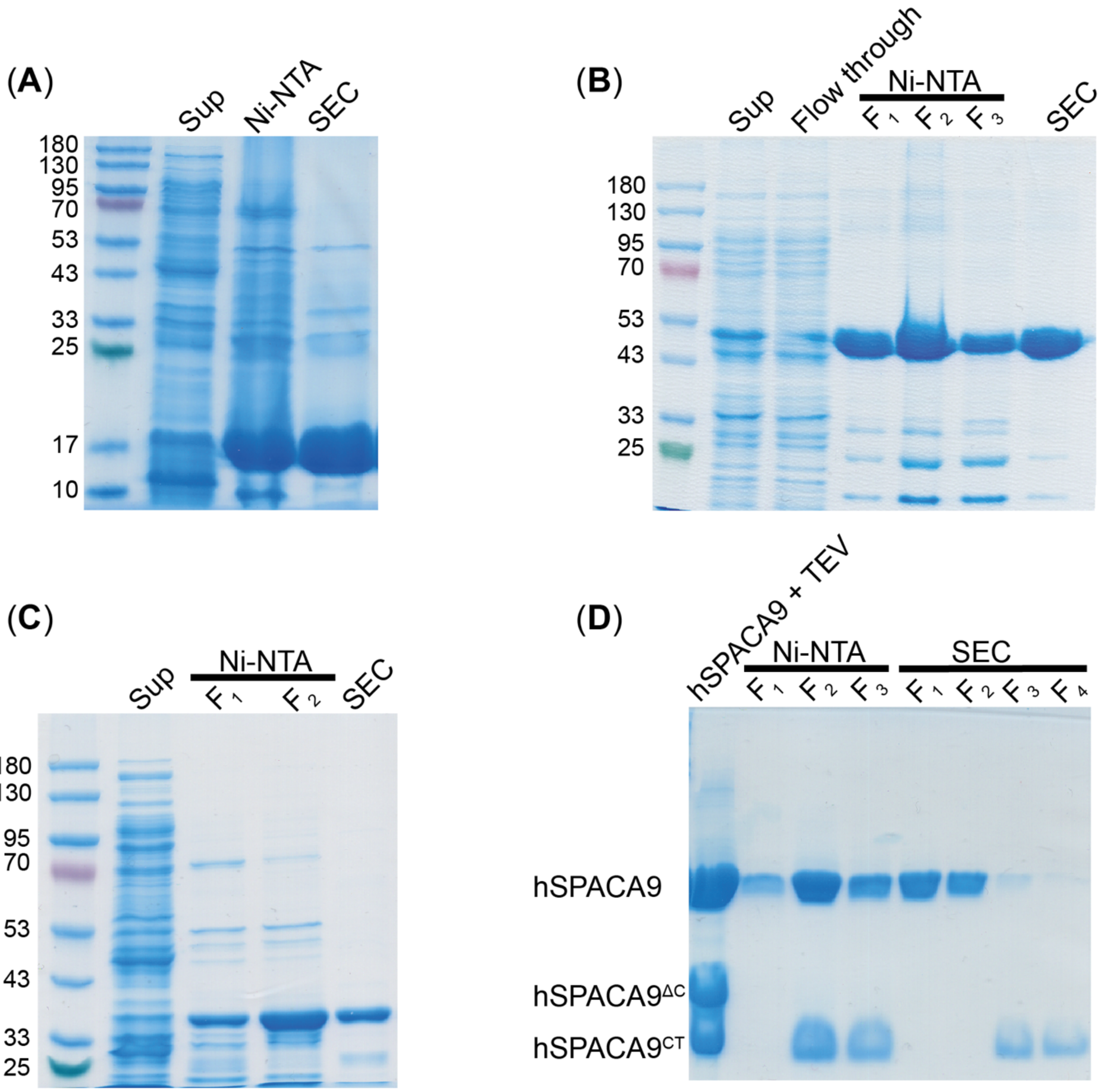


**Fig. S6. hSPACA9 constructs purification.** SDS-PAGE gel, stained with Coomassie blue, showing the expression and purification steps for (**A**) $hSPACA9^{\Delta C}$ and (**B**) $hSPACA9^{\Delta C\text{-}GFP}$. (**C**) $hSPACA9^{CT\text{-}GFP}$. Lanes from left to right: Clarified supernatant (Sup), Ni-column elution (Ni-NTA), and size exclusion chromatography (SEC) elution. (**D**) Following purification, recombinant hSPACA9 was subjected to TEV protease cleavage, generating three species: uncleaved full-length hSPACA9 and two cleavage products, $hSPACA9^{\Delta C}$ and $hSPACA9^{CT}$. The reaction mixture was then applied to a second Ni-NTA chromatography step to isolate His-tagged $hSPACA9^{CT}$, followed by final purification by SEC.

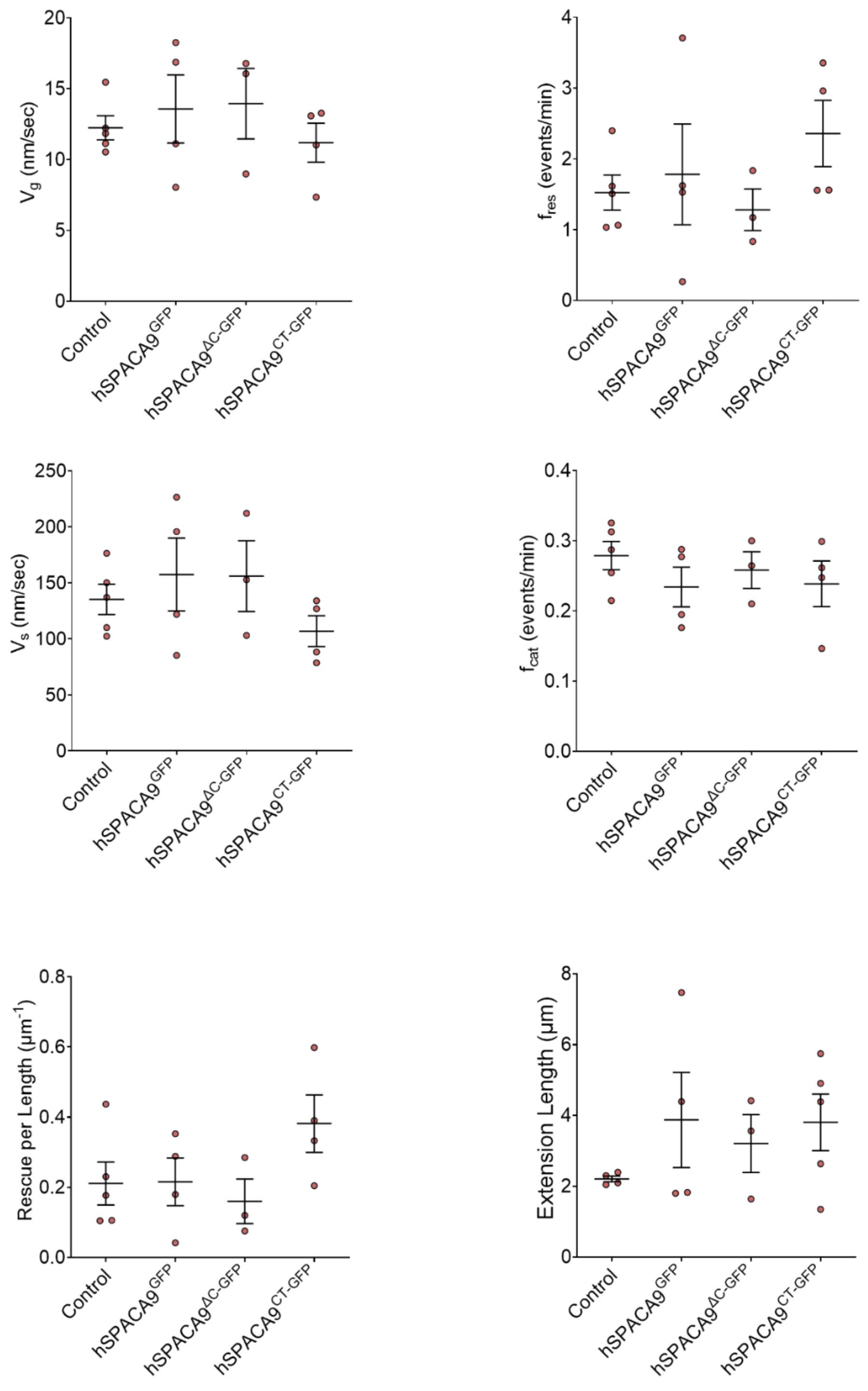


**Fig. S7. Quantification of microtubule dynamic properties of hSPACA9-GFP constructs**. Dot plots showing the effect of hSPACA9-GFP constructs [300 nM] compared with control conditions lacking hSPACA9 on microtubule growth rate ($V_g$), shrinkage rate ($V_s$), catastrophe frequency ($f_{cat}$), rescue frequency ($f_{res}$), rescue per length and extension length. Black lines show the overall mean and SE across replicates. All experiments were performed in the presence of 10 μM free tubulin.

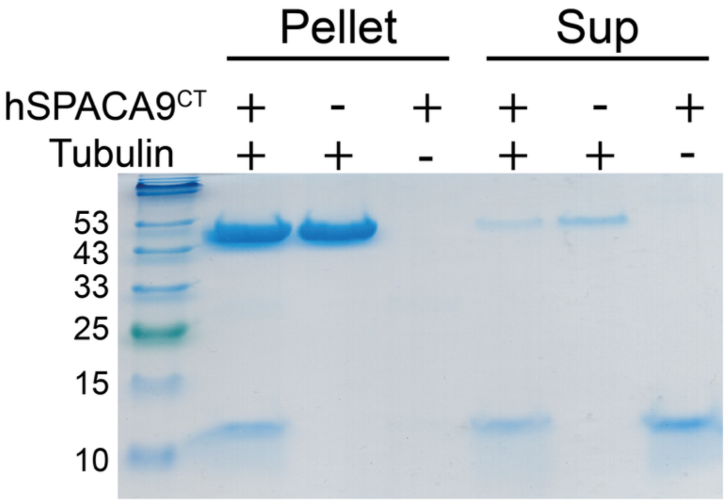


**Fig. S8. Co-sedimentation assay of hSPACA9$^{CT}$ with microtubules.** Purified hSPACA9$^{CT}$ was incubated with polymerizing microtubules and subjected to ultracentrifugation to separate microtubule-bound proteins (pellet) from unbound proteins (supernatant). Samples containing hSPACA9$^{CT}$ alone, tubulin alone, or both proteins were analyzed by SDS–PAGE followed by Coomassie staining. In the presence of polymerized microtubules, hSPACA9$^{CT}$ is detected in the pellet fraction together with tubulin, indicating association with the microtubule lattice. In contrast, hSPACA9$^{CT}$ alone remains predominantly in the supernatant fraction.

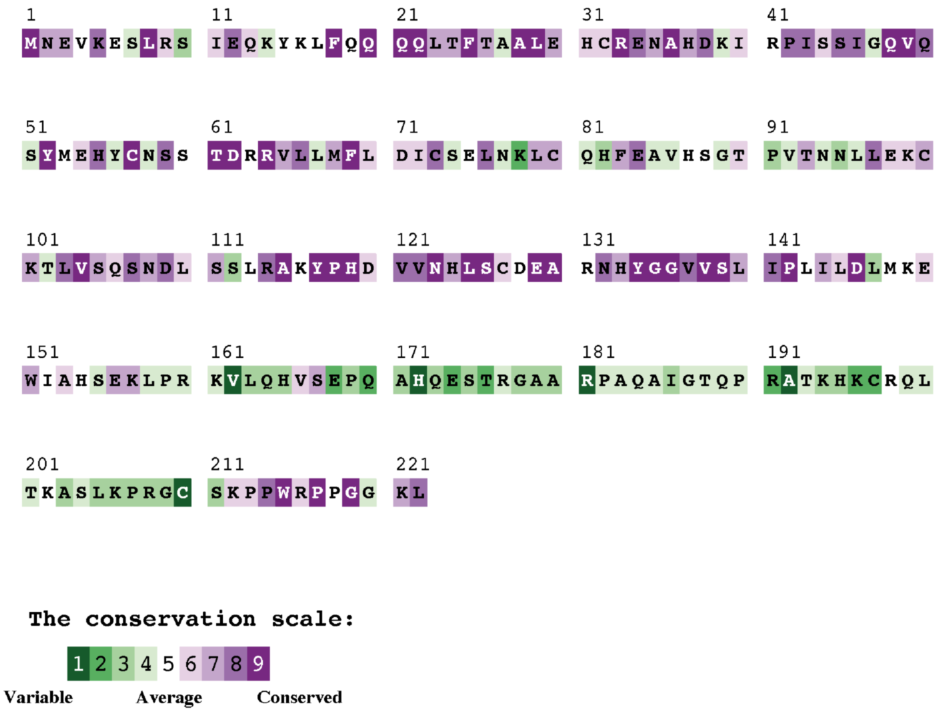


**Fig. S9. Conservation profile of full-length hSPACA9.** Residues are color-coded according to their evolutionary conservation scores, ranging from variable (green) to highly conserved (purple), as calculated by ConSurf. The profile highlights distinct conserved regions along the hSPACA9 sequence, including in the C-terminal tail (166-222), which is absent from the structural model but included in the sequence-based analysis.
